# Tumor-Intrinsic IL-17 Signaling Correlates with Multimodal Resistance Phenotypes Following Oncolytic Adenovirus Challenge

**DOI:** 10.64898/2026.03.27.714871

**Authors:** Eslam E. Abd El-Fattah, Mohamed Hammad

## Abstract

Oncolytic adenovirus (ADV) therapy faces heterogeneous responses, implying tumor-intrinsic resistance. We identify interleukin-17 (IL-17) signaling as a novel potential barrier associated with multi-modal cellular reprogramming. Transcriptomic analysis of ADV-treated 4T1 murine mammary carcinoma cells revealed specific upregulation of *Il17rb*, *Il17rd*, and *Il17f*, indicating viral induction of this inflammatory axis. The IL-17 signature correlates strongly with a cancer stemness phenotype. Metabolically, it associates with increased lipid metabolism and suppressed glycolysis, suggesting a state resistant to viral replication. Furthermore, it broadly negatively correlates with programmed cell death pathways (apoptosis, necrosis) while positively associating with pro-survival autophagy. IL-17 component expression effectively stratifies samples into distinct metastatic risk categories, underscoring its prognostic potential. Our findings reveal a previously unrecognized, tumor-intrinsic role for IL-17 signaling in ADV resistance, associated with enhanced stemness, altered metabolism, and impaired cell death. This nominates the IL-17 pathway as both a predictive biomarker and a therapeutic target for combination strategies.

**Highlights:**

- Oncolytic adenovirus infection selectively upregulates IL-17 receptor subunits (IL17RB, IL17RD) and IL17F ligand in 4T1 tumor cells
- IL-17 receptor expression strongly correlates with cancer stemness gene signatures, particularly through IL17RB and IL17RD
- The IL-17 axis associates with broad suppression of lytic cell death pathways (apoptosis, necrosis, necroptosis) while positively correlating with autophagy
- IL-17 pathway activity correlates with metabolic reprogramming favoring lipid turnover over glycolysis
- IL-17 expression levels stratify samples into distinct metastatic risk categories, suggesting biomarker potential

## 1. Introduction

Oncolytic adenoviruses are promising cancer therapies that leverage viral replication to lyse tumor cells and stimulate direct anti-tumor immunity. Their efficacy derives from tumor-selective replication and the immunogenic cell death that activates systemic immune responses **[1–3]**. These engineered oncolytic adenoviruses are designed to preferentially replicate within tumor cells, exploiting common cancer-specific vulnerabilities such as defective interferon responses and aberrant cell cycle signaling **[4, 5]**.

Despite this potential and encouraging preclinical results, patient outcomes are highly heterogeneous. This variability underscores significant biological barriers, including potent tumor-intrinsic resistance mechanisms. Cancer cells can evade virotherapy by downregulating viral entry receptors, activating intracellular antiviral defenses, and enhancing pro-survival pathways, which collectively limit therapeutic efficacy **[6, 7]**.

Furthermore, the hostile tumor microenvironment—marked by hypoxia, poor vasculature, and immunosuppressive cells—can restrict viral spread and protect resistant cells. A key resistant population is cancer stem cells (CSCs), which exhibit inherent defenses like quiescence and enhanced DNA repair **[8–12]**.

In this landscape, the interleukin-17 (IL-17) pathway is recognized for its complex, dual role in cancer. Typically a driver of pro-inflammatory responses via neutrophil recruitment, IL-17 signaling can, paradoxically, either promote anti-tumor immunity or drive pro-tumorigenic processes. This duality makes it a critical but complex factor in therapeutic resistance **[13–18]**.

Conversely, the IL-17 pathway also drives tumor progression via direct, tumor-intrinsic mechanisms. It stimulates cancer cell proliferation, survival, migration, and induces epithelial-mesenchymal transition (EMT). Critically, IL-17 maintains a cancer stem-like state, conferring resistance to chemo-and radiotherapy. It also promotes pro-tumor metabolic shifts. Elevated IL-17 signaling in multiple solid tumors correlates with poor prognosis. While classically part of the antiviral host defense, this pro-survival, stemness-promoting role suggests it may be co-opted by tumor cells to resist oncolytic viruses **[19–25]**.

This raises a critical, unexplored question: within the unique scenario of a therapeutic viral infection of a tumor, could the IL-17 axis—potentially induced by the virus itself—function paradoxically as a tumor-protective mechanism, enhancing cancer cell fitness and creating a barrier to successful oncolysis?

We hypothesize that oncolytic adenovirus infection induces tumor-intrinsic IL-17 signaling, which correlates with therapy resistance. To test this, we analyzed the transcriptomic response of 4T1 mammary carcinoma cells to adenovirus challenge. We aimed to: (1) characterize global transcriptional changes, (2) determine specific IL-17 pathway modulation, (3) investigate correlations between IL-17 activity and cancer hallmarks like stemness and metabolism, and (4) assess IL-17’s prognostic value for metastatic risk. Our results reveal a novel link between IL-17 and oncolytic virotherapy resistance, providing a new mechanistic target for combination therapies.

## 2. Materials and Methods

### 2.1 Differential Expression Analysis

Raw RNA-seq count data (GSE271202) were imported into R version 4.5.1 and processed using DESeq2 version 1.44.0. Genes with missing identifiers were removed, and sample metadata were organized with condition (control/ADV) and batch variables. The dataset comprised 3 biological replicates per condition (n = 3 ADV-treated, n = 3 control). The adenovirus used in the original study was human adenovirus type 5. Human adenoviruses exhibit restricted productive replication in murine cells due to species-specific barriers; they retain the capacity to enter cells and express early genes, allowing assessment of the initial transcriptional response to viral challenge rather than productive infection. A DESeq Data Set was created with design ∼ condition, filtering genes with <10 total counts. The DESeq2 workflow applied median-of-ratios normalization, dispersion estimation, and negative binomial modeling. Differential expression between ADV and control was tested via Wald tests (contrast: c(“condition”, “ADV”, “control”)), with significance defined as Benjamini–Hochberg adjusted p-value < 0.05.

### 2.2 Downstream Processing and Visualization

Significant genes (padj < 0.05) were sorted by ascending padj for downstream analysis. Visualization was performed using ggplot2 v4.0.1 and ggrepel v0.9.6. Three complementary plots were generated: (1) A volcano plot showing −log padj versus log FC, with genes meeting |log FC| > 1 and padj < 0.05 highlighted and the top 10 most significant genes labeled; (2) A bar plot summarizing counts of upregulated and downregulated genes; and (3) A horizontal bar plot of the top 20 significant genes, where bar length represents−log padj and color indicates log FC direction and magnitude. All figures were prepared in publication-quality format.

### 2.3 Comparative Analysis of IL-17 Receptor Expression and Functional Associations

All correlation analyses described below were performed across all samples pooled (ADV-treated and control; total n = 6), unless otherwise specified. This approach was employed to maximize statistical power and to identify associations between IL-17 pathway activity and cellular phenotypes that are consistent across experimental conditions.

#### 2.3.1 IL-17 Receptor Expression Across Experimental Conditions

Expression levels of IL-17 receptor genes (IL17RA, IL17RD, IL17F, and IL17RB) were compared between adenovirus (ADV)-treated and control (Ctrl) groups. Mean expression values with standard error bars were calculated, with individual data points overlaid to visualize sample-level variation. Statistical significance of group differences was assessed using two-sample t-tests, with results visualized in a multi-panel plot. Significance thresholds were denoted as: ***p < 0.001, **p < 0.01, *p < 0.05, and ns (not significant).

#### 2.3.2 Cancer Stemness Quantification and IL-17 Correlation Analysis

A composite cancer stemness score was derived from 32 established stemness marker genes. Expression values for each marker were z-score normalized across samples, and the overall stemness score was calculated as the mean of these normalized values. Pearson correlation analysis was performed between IL-17 receptor gene expression levels and stemness scores, with statistical significance defined at p < 0.05. Results were visualized to illustrate directional relationships and the strength of associations between IL-17 signaling and cancer stemness properties.

#### 2.3.3. IL-17 and Cell Death Pathway Analysis

Pathway definition: Seven cell death pathways (Apoptosis, Necrosis, Necroptosis, Autophagy, Ferroptosis, Cuproptosis, Pyroptosis) were defined with canonical gene sets including human/mouse orthologs. Gene sets were derived from established pathway databases [e.g. KEGG, Reactome, or literature-curated]. We made Activity scoring as Pathway scores calculated as mean z-score of available component genes per sample. Correlation analysis was made using Pearson correlations between IL-17 genes and pathway scores with p<0.05 significance threshold.

#### 2.3.4 IL-17 Regulation of Metabolic Pathways

The relationship between IL-17 signaling and cellular metabolism was assessed through systematic correlation analysis across predefined metabolic pathways. Pre-calculated correlation coefficients (Pearson or Spearman, range:-1 to 1) quantified relationships between specific IL-17-related genes (e.g., IL17A) and component genes within each metabolic pathway. All analyses were performed in R (version 4.5.1) using dplyr (version 1.1.4) for data manipulation and ggplot2 (version 4.0.1) for visualization.

Initial analysis employed a comparative heatmap visualizing all individual correlations using a blue-to-red gradient (negative to positive associations), with statistically significant correlations (p < 0.05) marked with asterisks. To derive pathway-level insights, a metabolic activation summary was calculated by aggregating significant findings: for each pathway, the mean correlation of all significant IL-17 gene associations served as a net metric of overall pathway activation/inhibition. This summary was presented as a bar chart indicating both the mean correlation and count of significant associations. The final integrated visualization combined detailed heatmap and pathway summary plots using the patchwork package for comprehensive presentation.

#### 2.3.5 Metastasis Pathway Integration and Risk Stratification

Metastasis-associated gene expression was quantified through pathway-specific scoring. For each sample, component genes of five distinct metastasis pathways (including EMT and Invasion signatures) were z-score normalized, and pathway scores were calculated as the mean normalized expression of respective gene subsets. Pearson correlation analysis assessed relationships between IL-17 family gene expression and individual metastasis pathway scores, with results visualized in a correlation heatmap highlighting significant associations (p < 0.05).

To evaluate the predictive relationship between IL-17 signaling and metastatic potential, a multiple linear regression model was constructed with the overall metastasis score as the dependent variable and all available IL-17 gene expression levels as independent variables. Furthermore, samples were stratified into four risk categories (Low, Medium-Low, Medium-High, High) based on quartiles of mean IL-17 pathway expression. Metastasis score distributions across these risk groups were visualized using violin plots and compared statistically using the Kruskal-Wallis test.

## 3. Results

### 3.1. Adenovirus Challenge Induces Global Transcriptional Reprogramming with Selective Upregulation of IL-17 Pathway and Its Subsequent Effect on Cancer Stemness

#### 3.1.1 Global Transcriptional Response

Differential gene expression (DGE) analysis identified 2,210 significantly altered transcripts (adjusted p<0.05, log 2-FC> 1), with the regulatory effect being nearly balanced quantitatively (1136 downregulated vs. 1074 upregulated) **(Figure 1A)**. However, the volcano plot **(Figure 1B)** revealed a profound asymmetry: while the upregulation range was confined to log 2-FC of approximately +4, the downregulatory response was significantly stronger in both magnitude and statistical significance, extending to log 2-FC of approximately-12 and a-log10 adjusted P value of approximately 150 **(Figure 1B)**. Genes like *Lox5*, *Sce2*, and *Mad* exemplify these extreme suppressive effects. Targeted analysis showed Myo10 as the most statistically significant transcript overall. Despite this, Lypd5 exhibited the greatest magnitude of downregulation, while transcripts like *Ltbp2*, *Ccnd1*, and *Axl* represented the most notable upregulated transcripts, confirming a complex, but suppressive-dominant, transcriptional signature **(Figure 1C)**.

**Figure 1:**
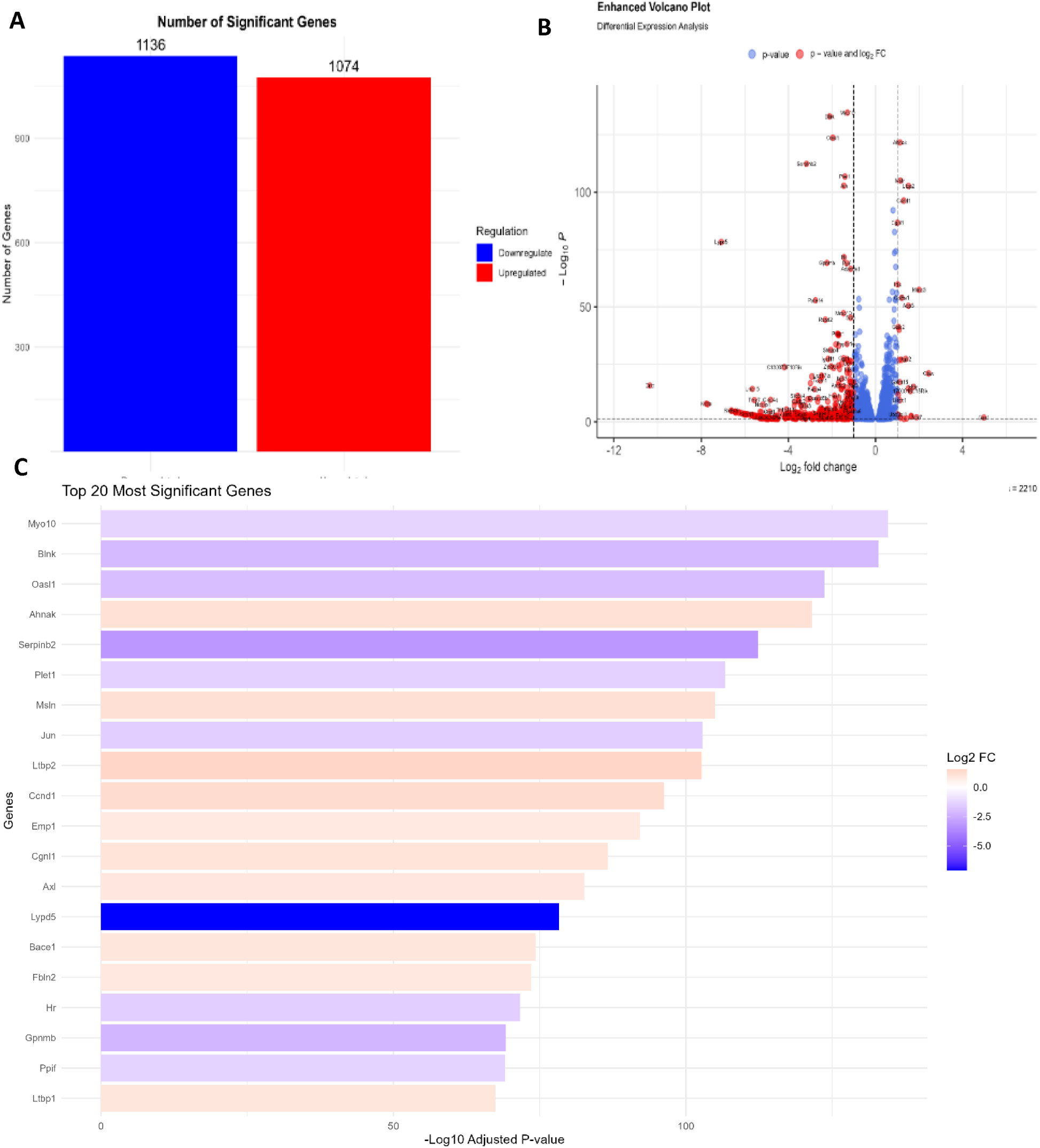
Global Differential Gene Expression Analysis Following Adenovirus Challenge. (A) Quantification of differentially expressed genes (DEGs). Bar plot summarizing the total number of genes identified as significantly upregulated (red, n=1,074) or downregulated (blue, n=1,136). **(B) Volcano plot** illustrating differential gene expression (DGE) analysis between ADV-challenged and Control 4T1 cells. Genes are plotted based on statistical significance (-log□□ adjusted p-value) and magnitude of expression change (log□ fold change). Red points indicate genes meeting both significance (adjusted p < 0.05) and fold-change (|log□FC| > 1) criteria; blue points indicate genes significant by p-value only. Dashed lines represent defined thresholds. **(C) Top 20 Most Significant Differentially Expressed Genes.** Bar length reflects-log□□ adjusted p-value; color indicates magnitude and direction of expression change (red/orange: upregulation; purple/blue: downregulation).

#### 3.1.2 Effect of IL-17 Pathway on cancer stemness

Among the inflammatory pathways represented in this transcriptional signature, the IL-17 axis emerged as particularly notable given its established roles in both antiviral immunity and tumor progression. We therefore specifically interrogated the expression of IL-17 family members. The differential expression analysis of the IL-17 family highlighted a specific transcriptional modulation of IL-17 pathway components induced by adenovirus (ADV) treatment in 4T1 cells. We observed a significant upregulation of three key elements—the ligand Il17f and the receptor subunits Il17rd and Il17rb **(Figure 2)**. This enhanced capacity for signal reception (IL-17RB, IL-17RD) and cytokine production (IL-17F) in treated tumor cells indicates virus-induced transcriptional priming of this pathway.

**Figure 2:**
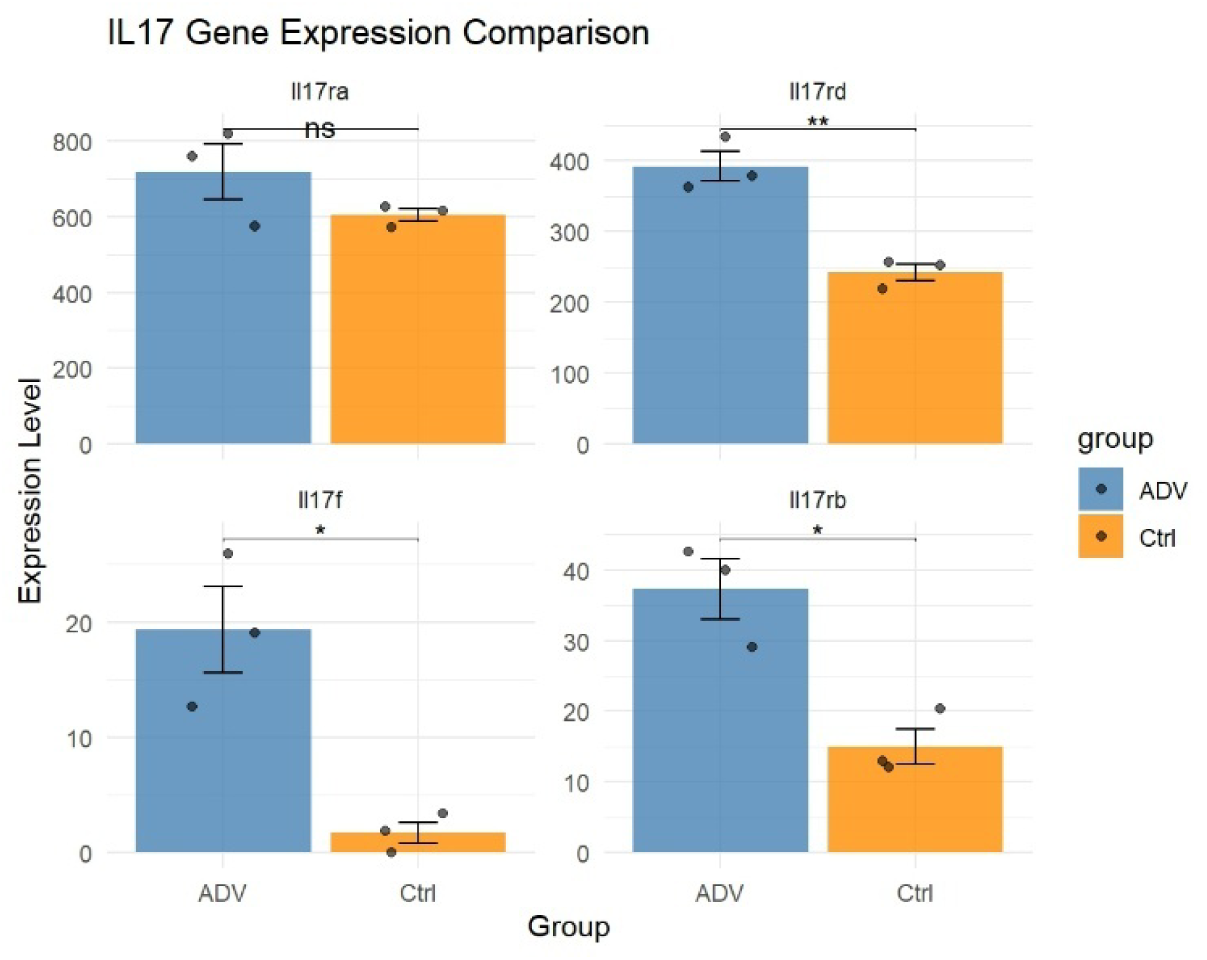
Selective Upregulation of IL-17 Pathway Components by Adenovirus Challenge. Bar plots display expression levels of Il17ra, Il17rd, Il17f, and Il17rb in ADV-challenged (blue) versus control (orange) samples. Data represent mean ± SEM with individual data points overlaid. Statistical significance assessed by two-sample t-test: *p < 0.05, **p < 0.01, ns = not significant.

### 3.2. IL-17 Axis Broadly Suppresses Lytic Cell Death Pathways While Promoting Autophagy

Given that oncolytic viruses exert their therapeutic effect through tumor cell lysis, we first examined whether the IL-17 signature correlates with cell death pathway activity—the most direct determinant of oncolytic efficacy. Correlation analysis focusing solely on cellular fate pathways revealed a generalized negative association of the IL-17 axis with lytic and programmed cell death mechanisms. Significant negative Mean Correlations were uniformly observed for Necroptosis, Apoptosis, and Necrosis, indicating that increased IL-17 gene expression strongly correlates with reduced activity of these critical cell death programs **(Figure 3)**. This strong, unidirectional negative association suggests that the expression of the IL-17 receptor components is associated with reduced expression of the core pathways responsible for executing these specific cell death programs.

**Figure 3:**
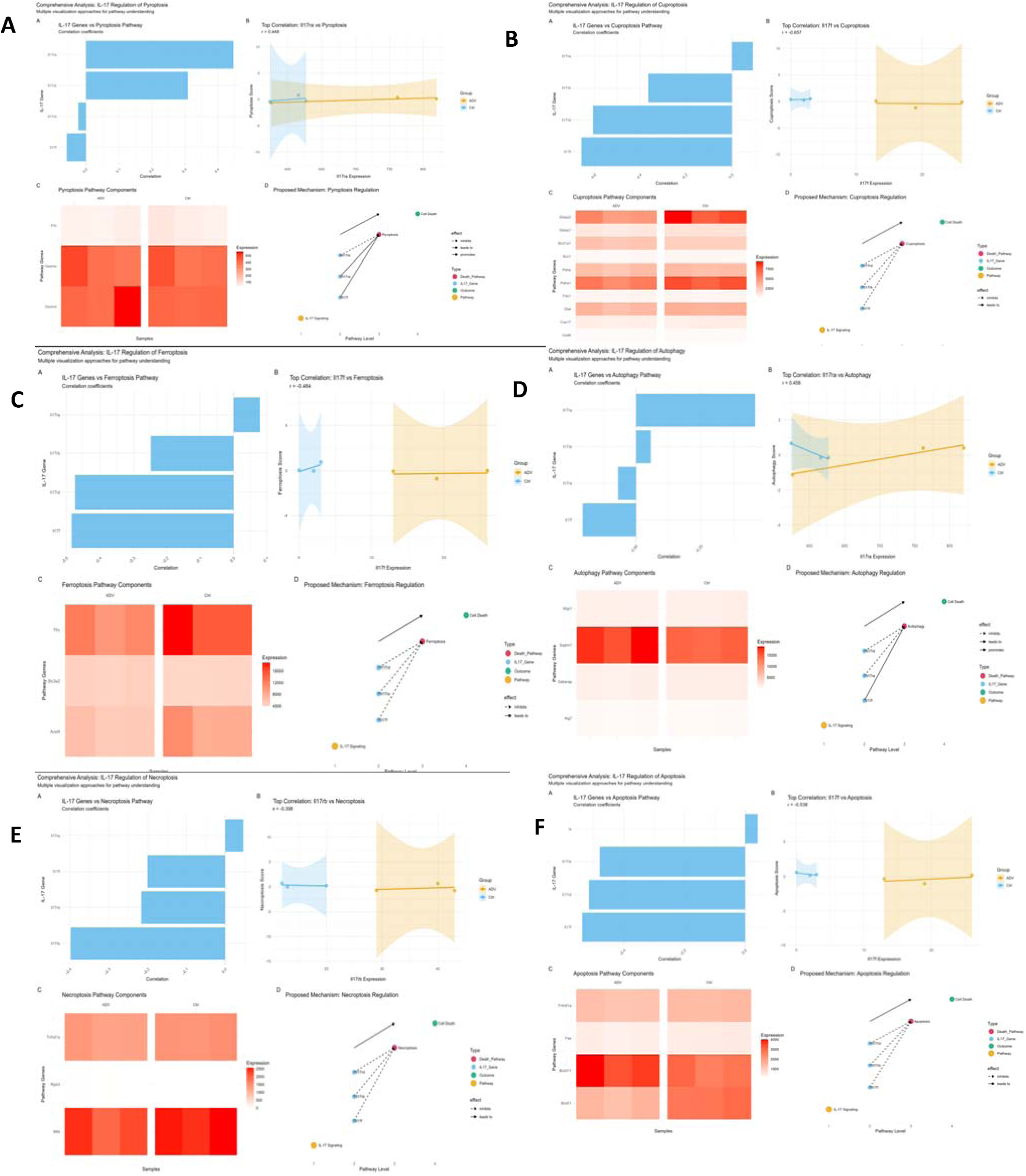
IL-17 Axis Correlates with Suppression of Lytic Cell Death Pathways. This composite figure displays correlations between IL-17 family gene expression and cellular fate pathways. A) proptosis, B) Cuproptosis. C) Ferroptosis, D) autophagy, E) Necroptosis, and F) Apoptosis. Each image has 4 subfigures (A-D), Individual pathway analyses. Each panel includes: (Left) Summary bar plot displaying mean Pearson correlation coefficient across significant IL-17 gene associations (orange: positive correlation; blue: negative correlation); (Right) Heatmap visualizing individual correlation coefficients between each IL-17 gene and pathway component genes (red: positive; blue: negative; *p < 0.05). (E) Proposed model: IL-17 pathway activity associates with suppression of lytic death programs (apoptosis, necrosis, necroptosis) while positively correlating with pro-survival autophagy.

This widespread negative correlation of cell death was specifically contrasted by a positive Mean Correlation observed for Autophagy. As Autophagy is a crucial cellular survival and resource recycling mechanism, its positive association with IL-17, coupled with the suppression of lytic cell death (Necroptosis and Necrosis) and controlled programmed death (Apoptosis), suggests that the IL-17 axis is associated with a robust cellular survival phenotype and thus encourages cancer cell survival within the treated 4T1 cells. Therefore, the regulatory signature of the IL-17 pathway appears to correlate with cell fate decisions, with negative associations to death execution programs and positive associations to survival-promoting autophagy.

### 3.3. IL-17 Receptor Expression Correlates Strongly with Cancer Stemness Signatures

The observed negative correlation between IL-17 pathway activity and cell death raises the question of which tumor cell subpopulations exhibit this death-resistant phenotype. Cancer stem cells (CSCs) represent intrinsically therapy-resistant subpopulations characterized by enhanced survival mechanisms. We therefore examined correlations between IL-17 pathway expression and a comprehensive stemness signature. To investigate the functional relevance of the IL-17 family expression profile, Pearson correlation analysis was performed against a calculated cancer stemness score **(Figure 4)**. We evaluated cancer stemness using multiple dimensions of stemness. For Core pluripotency factors, we evaluated Pou5f1, Sox2, Nanog, Klf4, and Myc and we evaluated embryonic stem cell markers using Fgf4, Utf1, Dppa2, Dppa4, and Dppa5 and cell surface stemness markers such as Prom1, Itga6, Itgb1, Cd44, and EPCAM. For signaling pathway components, we evaluated Bmi1, Notch1, Notch2, Hes1, Hey1, Gli1, Gli2, Smoothened, Patched1, Wnt1, Wnt3a, Beta-catenin, and Axin2. For EMT and stemness, we evaluated Vimentin, Snai1, Snai2, Twist1, and Zeb1. For Drug resistance, we evaluated Abcb1 and Abcg2. For Quiescence markers, we evaluated Nestin and Nes. We also evaluated cancer stem cell markers such as ALDH1a1, ALDH1a3, CD133, CD24, and CD34.

**Figure 4:**
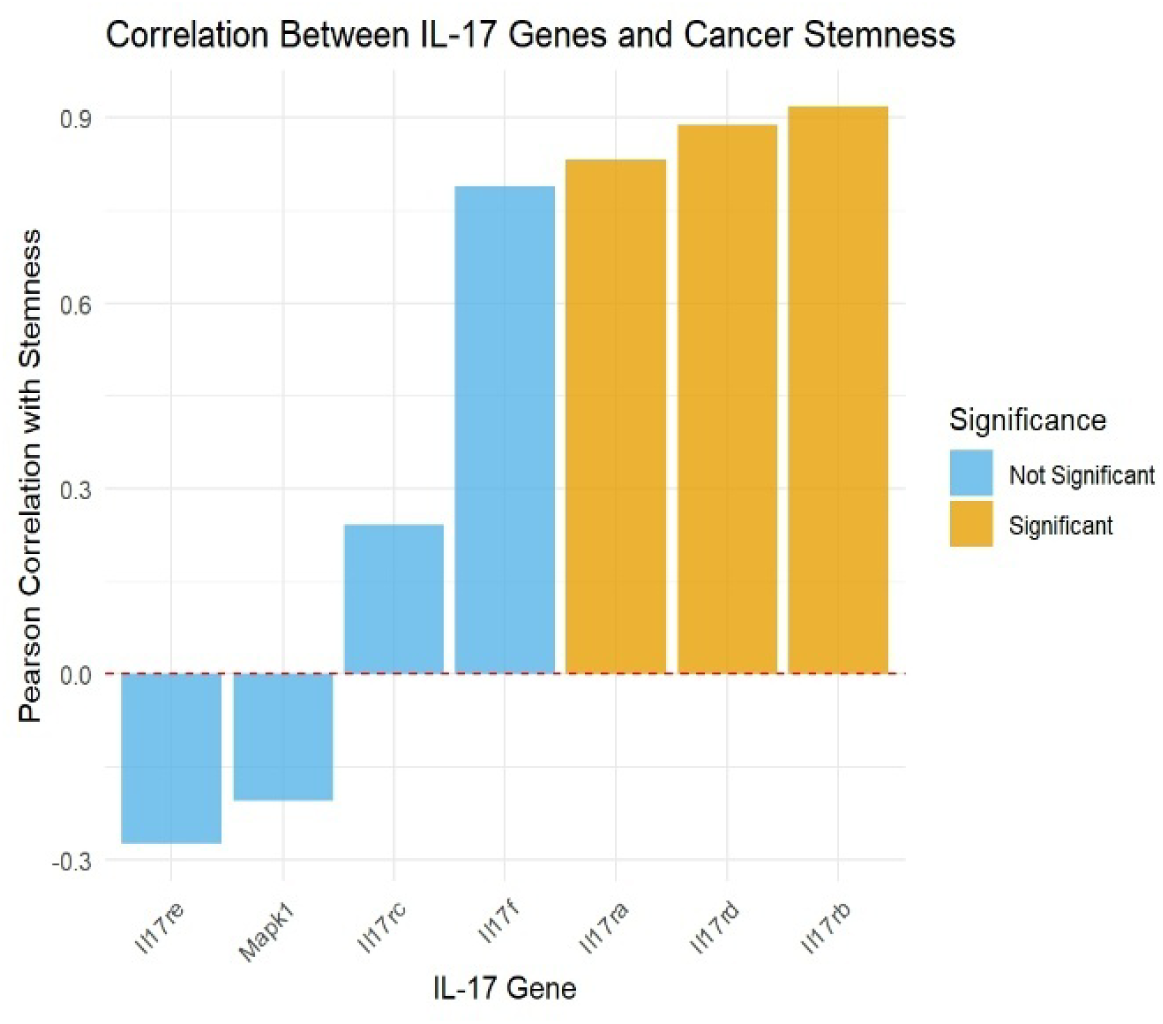
IL-17 Receptor Expression Correlates with Cancer Stemness. Bar plot depicting Pearson correlation coefficients between IL-17 family gene expression and composite stemness score derived from 32 canonical stemness markers. Orange bars indicate positive correlations; blue bars indicate negative correlations. Solid bars: p < 0.05; hatched bars: p ≥ 0.05 (not significant). Red dashed line indicates zero correlation.

The dominant finding was a strong, statistically significant positive correlation between key IL-17 genes and stemness. Specifically, Il17rb, Il17rd, and Il17ra all exhibited significant associations with increased stemness, suggesting that higher expression of these receptor components strongly correlates with a more aggressive, stem-like phenotype. Conversely, Il17re and Mapk1 showed negative correlations (approximately-0.30 and-0.20, respectively); however, these associations, along with the correlations for Il17rc and Il17f, were not statistically significant (blue bars). This data indicates that the receptor components Il17rb, Il17rd, and Il17ra are the primary IL-17 family members associated with the IL-17 axis to cancer stemness in this model. Notably, the same receptor subunits showing virus-induced upregulation (Il17rb and Il17rd) were among those most strongly correlated with stemness, suggesting that ADV challenge may prime tumor cells toward a stem-like, therapy-resistant phenotype via the IL-17 axis.

### 3.4. Metabolic Reprogramming by the IL-17 Axis Favors Lipid Metabolism Over Glycolysis

Maintenance of the stem-like, death-resistant phenotype requires metabolic adaptation. Cancer stem cells characteristically exhibit altered metabolic profiles distinct from bulk tumor cells, often favoring oxidative metabolism and lipid utilization over glycolysis. We therefore investigated whether IL-17 axis activity correlates with metabolic pathway alterations that could support such a resistant phenotype. The comparative correlation analysis **(Figure 5)** revealed a strong, bidirectional regulatory effect of the IL-17 gene family, particularly highlighting a net positive effect on lipid metabolism. Both Lipogenesis and Lipolysis exhibited the highest positive Mean Correlation values among all pathways studied, indicating a robust association between IL-17 expression and lipid turnover. This strong positive association was evidenced in the heatmaps by dense clusters of high correlation coefficients (deep red colors), primarily driven by the expression of *Il17b* and *Il17a* genes within the components of the lipogenesis and lipolysis pathways.

**Figure 5:**
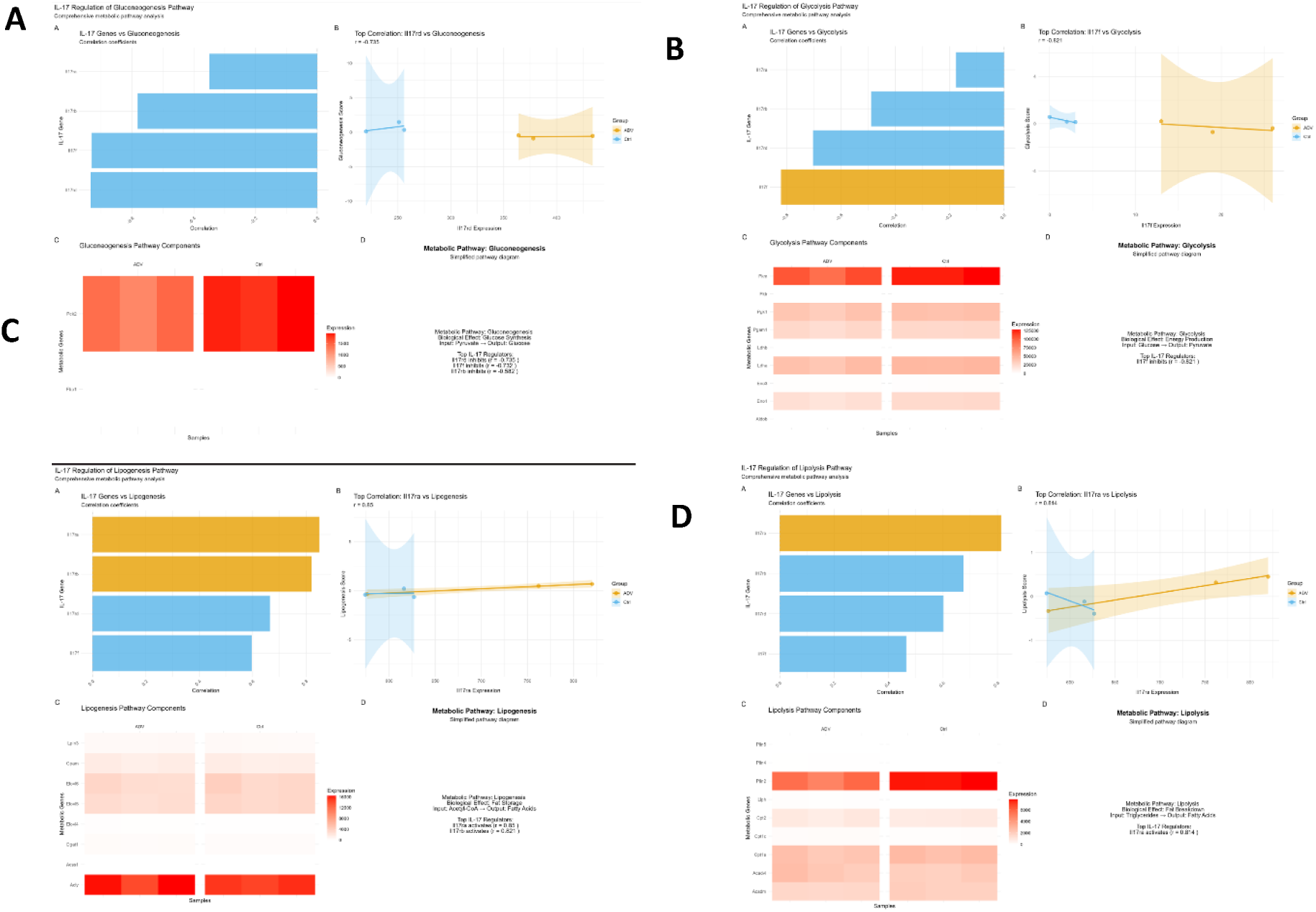
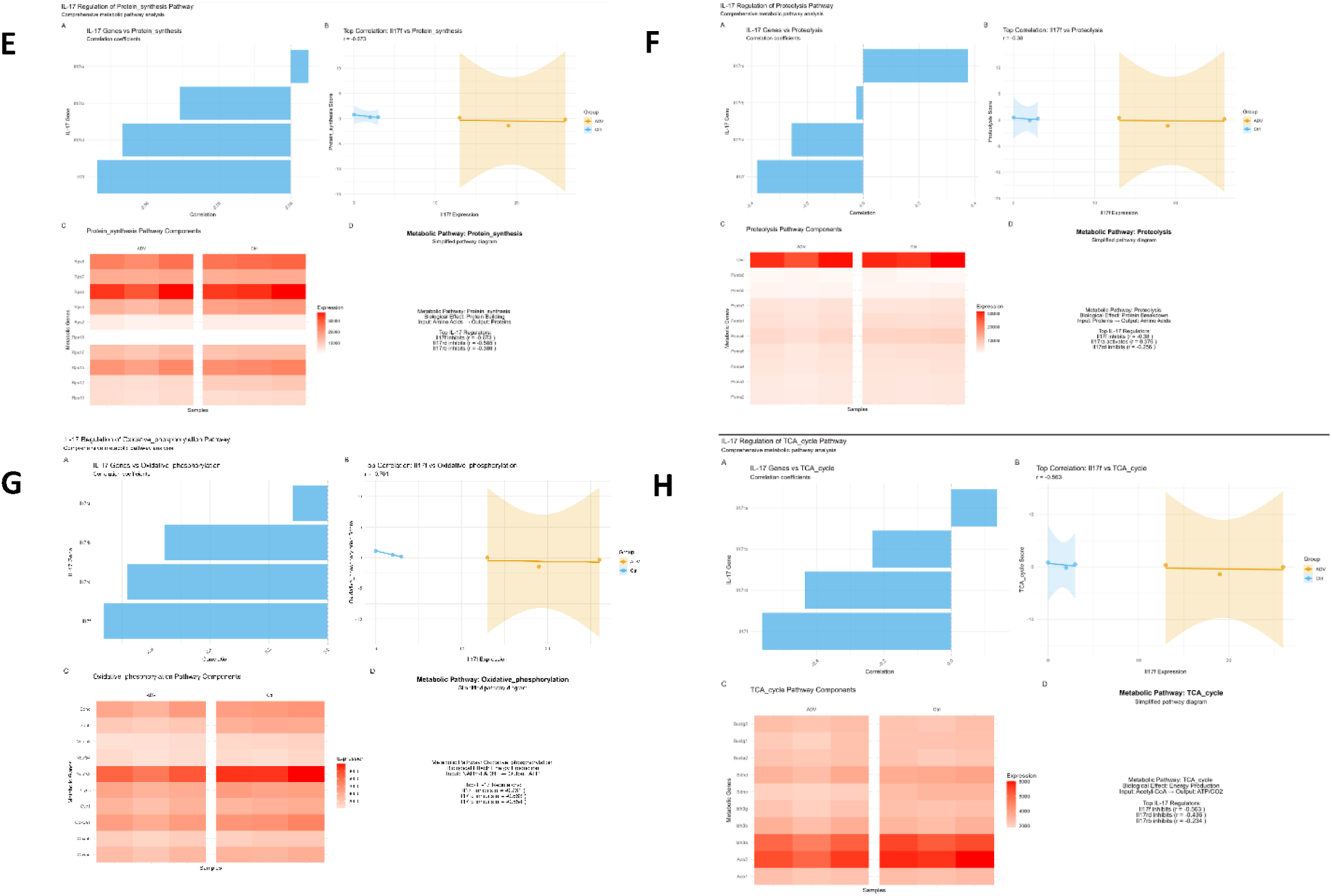
Comparative Correlation Analysis of IL-17 Gene Expression and Metabolic Pathway Components. This composite figure displays the statistical relationship between the expression of IL-17 family genes and the gene components of various metabolic pathways, including **A) Gluconeogenesis, B) Glycolysis, C) Lipogenesis, D) Lipolysis, E) protein synthesis, F) Proteolysis, G) Oxidative Phosphorylation, H) TCA cycle.** Each primary pathway analysis (e.g., Panels A-B, C-D, etc.) includes three key components: Summary Bar Plot (A, C, E, G, I, K, M, O): Displays the Mean Correlation calculated across all statistically significant IL-17 gene associations for that pathway. This metric represents the net regulatory effect: positive mean correlation (yellow/orange) indicates net activation, and negative mean correlation (blue) indicates net inhibition. Top Correlation Plot (B, D, F, H, J, L, N, P): Illustrates the specific Pearson correlation between the top correlated IL-17 gene and a key component of the metabolic pathway. Heatmap: Visualizes the individual Pearson correlation coefficients between each IL-17 Gene and each Metabolic Pathway component. Color intensity and direction (red: positive correlation; blue: negativ correlation) reflect strength, with statistically significant associations (*p < 0.05) highlighted or numerically marked.

Conversely, the data indicated a significant, specific negative correlation with Glycolysis, which displayed the largest negative Mean Correlation value. This strong negative effect was driven by specific subunits, notably *Il17f* and *Il17d*, which displayed strong negative correlations (deep blue colors) with glycolytic markers (e.g., *Il17f* correlation approaching r =-0.82*). Furthermore, pathways such as the TCA cycle, Gluconeogenesis, Protein Synthesis, and Proteolysis did not yield a critical mass of statistically significant associations to establish a definitive net regulatory trend. While isolated strong numerical correlations were noted (e.g., *Il17d* correlation of r = - 0.68 in Protein Synthesis), the absence of a clear net Mean Correlation suggests the metabolic association of the IL-17 family is highly specific, primarily focused on modulating lipid synthesis/breakdown and negative association with glucose catabolism. This metabolic signature enhanced lipid turnover coupled with reduced glycolytic gene expression is consistent with the metabolic profile of cancer stem cells and therapy-resistant tumor subpopulations, which often rely on fatty acid oxidation rather than glycolysis for energy production.

### 3.6. Summary of Correlational Findings

The analysis highlights strong, bidirectional regulatory effects of the IL-17 gene family on cellular metabolism. Specifically, the IL-17 axis exerts a robust positive regulation on lipid metabolism. As illustrated in the heatmap **(Figure 6A)**, *IL17b* and *IL17a* drive a dense cluster of strong positive correlations within the lipogenesis and lipolysis pathways. This is corroborated by the summary analysis **(Figure 6B)**, where both lipogenesis (n=2 significant genes) and lipolysis (n=1) exhibit the highest positive mean correlation values, indicating a net activating effect on lipid turnover. Conversely, the data indicates significant inhibition of glycolysis. *IL17f* and *IL17d* display strong negative correlations with glycolytic markers (e.g., *IL17f*: r =-0.82*), resulting in the largest negative mean correlation observed (n=1). While isolated strong numerical correlations were present in other pathways—such as *IL17d* in protein synthesis (r=-0.68)—pathways including oxidative phosphorylation, gluconeogenesis, and the TCA cycle did not yield a critical mass of statistically significant associations (p < 0.05) to establish a definitiv net regulatory effect.

**Figure 6:**
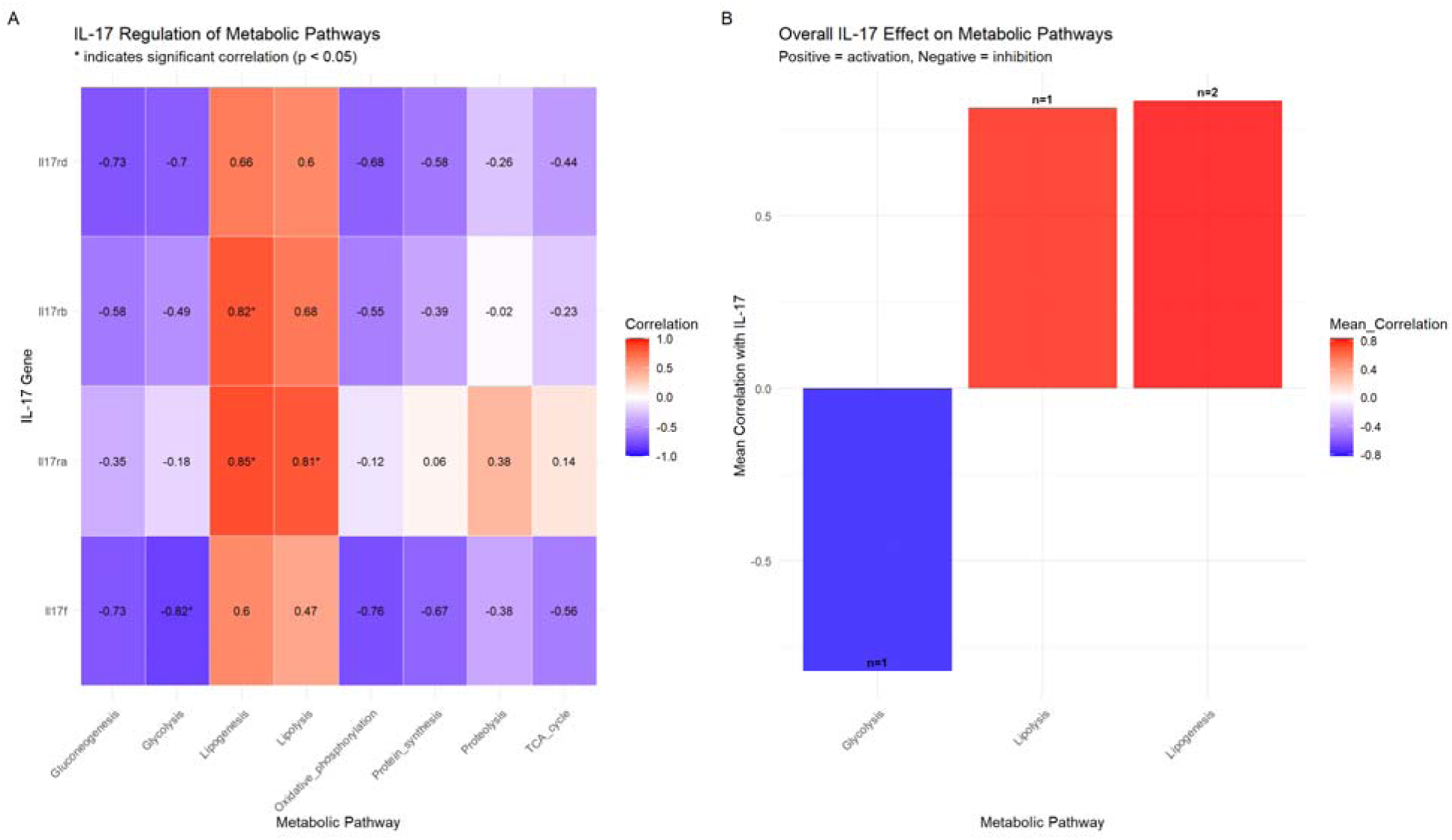
Summary of IL-17-Metabolic Pathway Correlations. (A) Heatmap summarizing correlations betwee IL-17 family genes (columns) and metabolic pathways (rows). Color scale represents Pearson correlation coefficient (red: positive; blue: negative); *p < 0.05. (B) Bar plot showing mean correlation for pathways with statistically significant associations. n indicates number of significant gene-level correlations per pathway.

### 3.7. IL-17 Expression Stratifies Samples by Metastatic Risk

Having established that IL-17 pathway activity correlates with multiple pro-survival phenotypes including death resistance, stemness, and metabolic adaptation, we evaluated whether this signature has prognostic value for identifying patients at elevated risk for aggressive disease.

We stratified patient samples based on IL-17 expression levels and their associated metastatic potential. Using a continuous Metastasis Pathway Score, samples were categorized into discrete risk groups: Low Risk (Score = 0.5), Medium-Low Risk (Score = 0.6), and combined Medium-High/High Risk (Score = 0.7). As shown in **Figure 7A**, a clear positive trend was observed between increasing IL-17 expression and escalation in metastatic risk category. This risk stratification was consistent across treatment conditions, with both the treatment (ADV) and control (Ctrl) groups showing the same pattern of risk distribution based on IL-17 expression. Although the overall comparison across risk categories did not reach statistical significance (p = 0.37), likely due to limited sample size, the consistent directionality of the trend across both ADV-treated and control conditions suggests a potential biological relationship that warrants validation in larger cohorts. These data suggest that IL-17 expression may have potential as a biomarker for stratifying metastatic risk, pending validation in appropriately powered studies.

**Figure 7:**
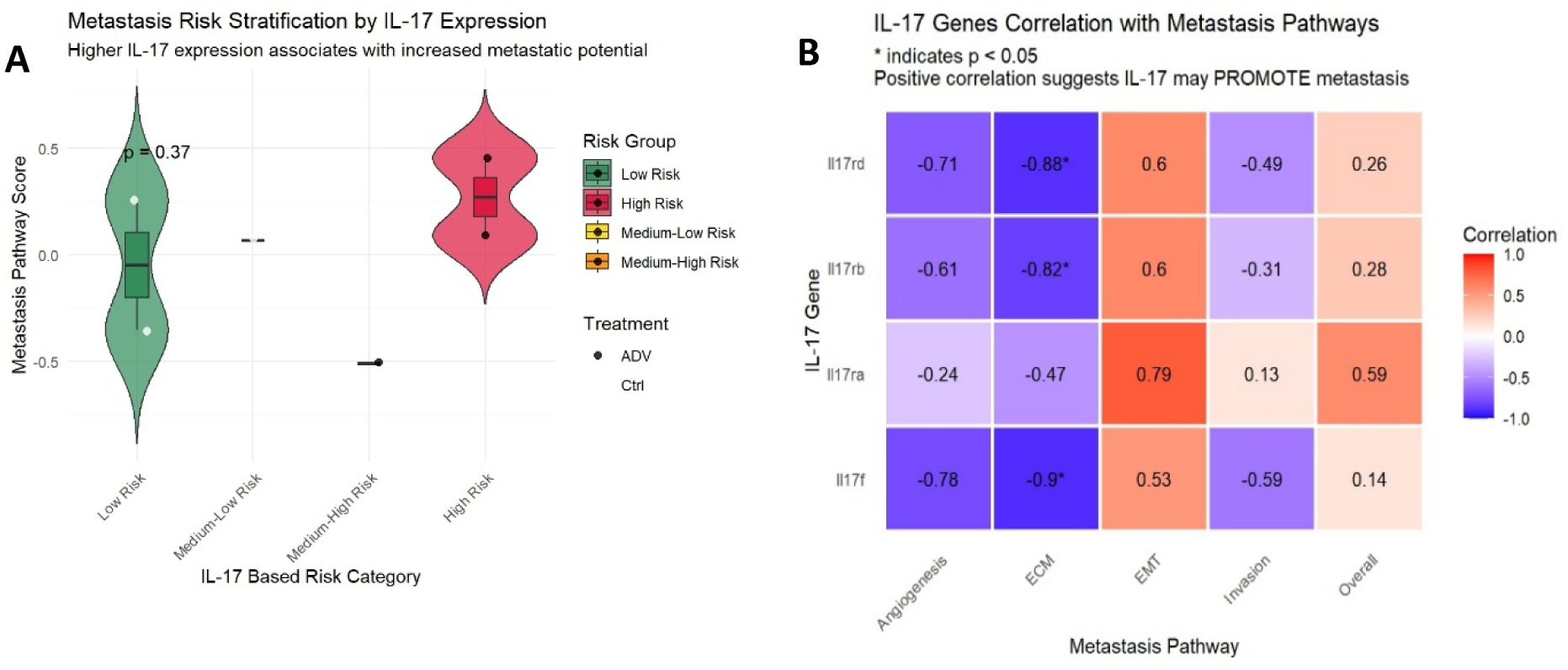
IL-17 Expression and Metastatic Risk Stratification. This composite figure summarizes the role of the IL-17 gene family in cancer metastasis. **(A)** Violin plot showing distribution of Metastasis Pathway Score across IL-17 expression-based risk categories (Low, Medium-Low, Medium-High, High). Individual points colored by experimental condition (ADV: blue; Control: orange). Kruskal-Wallis test p = 0.37 (not significant; see Limitations). **(B)** Heatmap displaying Pearson correlation coefficients between IL-17 family genes and five metastasis-associated pathway scores (Angiogenesis, ECM Remodeling, EMT, Invasion, Overall Metastasis). *p < 0.05.

The correlation heatmap **(Figure 7B)** depicts the relationship between the expression of IL-17 family genes (IL-17a, IL-17b, IL-17d, IL-17f) and five distinct metastasis-associated pathways. Several IL-17 isoforms exhibited significant negative correlations (p < 0.05, denoted by *) with specific pathways, most notably IL-17d, IL-17b, and IL-17f. The strongest negative correlations (approximately-0.9) were observed. In contrast, IL-17a demonstrated a moderate positive correlation (0.79) with one pathway and no significant negative associations. The overall pattern of negative correlations for IL-17b, d, and f suggests a complex, potentially regulatory role for these isoforms in the measured metastasis pathways, warranting further investigation into their specific mechanistic functions. To preliminarily assess the translational relevance of these findings, we note that IL-17 receptor expression has been reported in human breast cancer specimens and correlates with clinical outcomes 10.3389/fonc.2023.1171254. Future validation using The Cancer Genome Atlas (TCGA) breast cancer cohort or similar resources will be essential to confirm the prognostic utility of IL-17 pathway activity identified in this preclinical model.

Collectively, our analyses reveal that adenovirus-induced IL-17 pathway activation correlates with a coordinated phenotype encompassing: (1) suppression of lytic cell death pathways with concurrent autophagy activation; (2) enhanced expression of cancer stemness gene signatures; and (3) metabolic reprogramming favoring lipid turnover over glycolysis. This constellation of features is consistent with a therapy-resistant cellular state that may limit oncolytic virus efficacy.

## 4. Discussion

The variable efficacy of oncolytic adenoviruses (ADV) points to undefined tumor-intrinsic resistance. Our transcriptomic analysis of ADV-challenged 4T1 cells identifies IL-17 pathway activation as a novel correlate of resistance. The finding that ADV upregulates *Il17rb*, *Il17rd*, and *Il17f* indicates that tumors may co-opt this inflammatory axis to limit viral lysis, positioning IL-17 within a tumor-autonomous survival network. Correlation analyses link this IL-17 signature to a coordinated pro-survival phenotype, including enhanced stemness, metabolic reprogramming, and suppression of cell death pathways **[12, 26]**. The upregulation of *Il17rb* and *Il17rd* is mechanistically suggestive. IL-17RB pairs with IL-17RA, while IL-17RD is an atypical receptor known to modulate signaling pathways like FGF, often providing inhibitory feedback **[27]**.

Their co-induction alongside *Il17f* indicates a shift toward a signaling topology that may favor autocrine or paracrine loops within the tumor cell population **[28–30]**. This viral induction of a “self-signaling” inflammatory module could represent an evolutionary hijacking; by triggering a pathway associated with tissue repair and barrier defense, the tumor may be activating a program designed to maintain epithelial integrity and limit damage in this case, damage inflicted by the oncolytic virus. Our data showing strong positive correlations between Il17RB/RD/RA expression and a composite stemness score provide correlational validation for this hypothesis, linking the pathway directly to a therapy-resistant cellular state.

The connection between IL-17 signaling and cancer stemness is emerging as a critical axis in therapy resistance **[21]**. CSCs are characterized by enhanced DNA repair, quiescence, expression of drug efflux pumps, and resistance to apoptosis and oxidative stress a phenotype intrinsically resistant to both conventional therapies and viral oncolysis **[31, 32]**. Our finding that ADV-induced IL-17 receptor expression correlates with a stemness signature enriched for core pluripotency factors (*Pou5f1*, *Sox2*, *Nanog*), EMT drivers (*Snai1*, *Twist1*, *Zeb1*), and drug resistance mediators (*Abcb1*, *Abcg2*) suggests that the IL-17 axis may be associated with this refractory subpopulation. This aligns with recent studies showing that IL-17 can promote a stem-like phenotype in colorectal and pancreatic cancers by activating downstream effectors such as NF-κB and STAT3, which in turn transcriptionally upregulate stemness genes **[33, 34]**. Therefore, the IL-17 pathway may serve as a molecular node, associated with a transcriptional program that correlates with dedifferentiation of tumor cells toward a more primitive, resilient, and self-renewing state, thereby potentially representing a reservoir of cells impervious to ADV-mediated killing.

Perhaps the most striking and actionable finding of our study is the systematic metabolic reprogramming associated with the IL-17 signature. We identified a clear dichotomy: a strong, specific negative correlation with glycolysis driven primarily by *Il17f* and *Il17d*, coupled with a robust positive association with lipid metabolic pathways (lipogenesis and lipolysis) driven by *Il17b* and *Il17a*. This metabolic shift has profound implications for viral replication and oncolysis. Glycolysis is the primary source of energy (ATP), biosynthetic precursors (nucleotides, amino acids), and reducing equivalents (NADPH) necessary for the rapid, lytic replication cycle of adenovirus **[35, 36]**. By correlating with reduced expression and/or activity of glycolytic enzymes, the IL-17 axis may effectively be associated with metabolic conditions that limit “ the virus of the resources required for efficient replication, progeny production, and spread. Concurrently, the positive association with lipid metabolism may serve a dual pro-survival function. First, enhanced lipid synthesis provides membranes for organelle biogenesis and survival signaling platforms **[37]**. Second, increased lipolysis and fatty acid oxidation (β-oxidation) can generate ATP through mitochondrial respiration, a more efficient but slower process that supports long-term cell survival under stress, exactly the phenotype a CSC would adopt **[38, 39]**. This metabolic rewiring from a glycolytic, anabolic state to a lipid-catabolic, energy-efficient state mirrors the metabolic adaptations seen in quiescent, therapy-resistant cells and may represent a sophisticated, non-immunological defense strategy against the viral threat.

Complementing the metabolic signature, our analysis reveals that the IL-17 axis correlates with a profound recalibration of cellular fate decisions, broadly negatively associating with programmed lytic cell death while positively correlating with autophagy. The significant negative correlations with apoptosis, necrosis, and necroptosis indicate a reduced expression of cell death pathway genes. Adenoviruses themselves encode proteins (e.g., E1B-19K) to inhibit host apoptosis **[40, 41]**, but our data suggest the host cell may amplify this inhibition via IL-17 signaling, potentially through upregulation of anti-apoptotic BCL-2 family members or inhibitors of apoptosis (IAPs). The concurrent positive association with autophagy is particularly significant. Autophagy is a double-edged sword in viral infections; while it can degrade viral components (virophagy), it more often serves as a pro-viral process by recycling nutrients to support replication or as a cell survival mechanism during nutrient stress **[42]**. In our context, where glycolysis gene expression is reduced, the IL-17-associated increase of autophagy may function as a pro-survival, nutrient-salvaging pathway, allowing tumor cells to persist in a metabolically restricted state induced by both the virus and their own IL-17 response. This triad death pathway suppression, survival autophagy, and glycolytic reduction suggests a cell entering a state of “dormancy” or “persistence” to weather the viral challenge.

The translational potential of these mechanistic insights is immediately apparent in the risk stratification analysis. The finding that IL-17 expression levels may categorize samples into distinct metastatic risk groups, independent of ADV treatment, underscores its potential role as a fundamental determinant of tumor aggressiveness. This positions IL-17 pathway activity not merely as a resistance mechanism to a specific therapy but as a core component of a tumor’s intrinsic “fitness” program. From a clinical perspective, this suggests that pre-treatment assessment of IL-17 signaling (via gene expression, immunohistochemistry for receptors, or perhaps circulating IL-17 ligands) could identify patients whose tumors possess this built-in resistance network. These patients might be predicted to derive suboptimal benefit from oncolytic adenovirus monotherapy.

Based on our data, we propose a synergistic combination of oncolytic adenovirus and IL-17 pathway inhibitors. We hypothesize that blocking IL-17 will dismantle resistance by simultaneously sensitizing cancer stem cells to lysis, restoring glycolytic metabolism for viral replication, and reactivating apoptosis. Several classes of IL-17 pathway inhibitors are clinically available and have established safety profiles from their use in autoimmune diseases like psoriasis and psoriatic arthritis **[43]**. These include Monoclonal antibodies against IL-17A (Secukinumab, Ixekizumab) **[44]**, Monoclonal antibodies against the IL-17 receptor A (Brodalumab), which would block signaling from multiple IL-17 ligands and small molecule inhibitors targeting downstream signaling nodes **[45]** are in development as a combination therapy regimen could be envisioned where an IL-17 pathway inhibitor is administered before or concurrently with intratumoral ADV injection. The inhibitor could “prime” the tumor microenvironment by disabling the tumor-intrinsic resistance program we have identified, thereby creating a permissive state for enhanced viral replication, spread, and immunogenic cell death. This approach aligns with the growing paradigm of using targeted agents to modulate the tumor microenvironment and improve the efficacy of immunotherapies.

## 5. Conclusion

In conclusion, our work identifies a potential resistance mechanism wherein oncolytic adenovirus challenge paradoxically induces an IL-17-associated transcriptional program that correlates with molecular features of therapy resistance. This program is associated with a stem-like, metabolically adapted, and death-resistant cellular state. These insights transform IL-17 from a cytokine of interest in inflammation into a candidate therapeutic target in oncolytic virotherapy. The proposed combination of oncolytic adenovirus with IL-17 pathway blockade represents a rationally designed, key translatable strategy to overcome a potential barrier to efficacy. Future work should prioritize *in vivo* validation of this combination along with functional studies to establish causation, aiming to convert the heterogeneity of patient responses into consistent and durable therapeutic success, ultimately expanding the reach and impact of oncolytic virotherapy in the clinical armamentarium against cancer.

## 6. Resource availability

**Lead contact:**

Requests for further information and resources should be directed to and will be fulfilled by the lead contact, **Eslam E. Abd El-Fattah, MBA, PhD**

Department of Biochemistry, Faculty of Pharmacy, Delta University for Science and Technology, Gamasa, Egypt

Department of Developmental and Stem Cell Biology, Beckman Research Institute at City of Hope Comprehensive Cancer Center, Duarte, CA 91010, USA

Tel: +1-443-465-4055

### Materials availability

The RNA-seq data analyzed in this study are publicly available from the Gene Expression Omnibus (GEO) under accession number GSE271202.

## Data and code availability

Original data are publicly available and code for analysis is available upon request.

## 7. Acknowledgments

## Materials availability

This study did not generate new unique reagents.

## Author Contributions

E.A.E.: Conceptualization, Methodology, Formal Analysis, Writing – Original Draft. M.H.: Writing – Review & Editing, revision.

## References

1. Omolekan TO, Folahan JT, Tesfay MZ, Mohan H, Dutta O, Rahimian L, Ferdous KU, Ghavimi R, Cios A, Beng TK et al: Viral warfare: unleashing engineered oncolytic viruses to outsmart cancer’s defenses. 2025, Volume 16 - 2025.

2. Rivera-Orellana S, Bautista J, Palacios-Zavala D, Ojeda-Mosquera S, Altamirano-Colina A, Alcocer-Veintimilla M, Parrales-Rosales G, Izquierdo-Condoy JS, Vásconez-González J, Ortiz-Prado E et al: Oncolytic virotherapy and tumor microenvironment modulation. Clinical and experimental medicine 2025, 25(1):256.

3. Zabelina DS, Osipov ID, Maslov DE, Kovner AV, Vasikhovskaia VA, Demina DS, Romanov SE, Shishkina EV, Davydova J, Netesov SV et al: Ad6-Based GM-CSF Expressing Vector Displays Oncolytic and Immunostimulatory Effects in an Immunocompetent Syrian Hamster Model of Cholangiocarcinoma. 2025, 17(2):162.

4. Tan EW, Abd-Aziz N, Poh CL, Tan KO: Engineered Oncolytic Adenoviruses: An Emerging Approach for Cancer Therapy. Pathogens (Basel, Switzerland) 2022, 11(10).

5. Gupta A, Chavan SR, Gadepalli R, Pareek P: Oncolytic viruses in head and neck cancers: clinical applications and therapeutic potential. 2025, Volume 16 - 2025.

6. Bai R, Chen N, Li L, Du N, Bai L, Lv Z, Tian H, Cui J: Mechanisms of Cancer Resistance to Immunotherapy. Frontiers in oncology 2020, 10:1290.

7. Le Boeuf F, Bell JC: United virus: the oncolytic tag-team against cancer! Cytokine & growth factor reviews 2010, 21(2-3):205–211.

8. Kim SK, Cho SW: The Evasion Mechanisms of Cancer Immunity and Drug Intervention in the Tumor Microenvironment. Frontiers in pharmacology 2022, 13:868695.

9. Labani-Motlagh A, Ashja-Mahdavi M, Loskog A: The Tumor Microenvironment: A Milieu Hindering and Obstructing Antitumor Immune Responses. 2020, Volume 11 - 2020.

10. Phi LTH, Sari IN, Yang Y-G, Lee S-H, Jun N, Kim KS, Lee YK, Kwon HY: Cancer Stem Cells (CSCs) in Drug Resistance and their Therapeutic Implications in Cancer Treatment. 2018, 2018(1):5416923.

11. Phi LTH, Sari IN, Yang YG, Lee SH, Jun N, Kim KS, Lee YK, Kwon HY: Cancer Stem Cells (CSCs) in Drug Resistance and their Therapeutic Implications in Cancer Treatment. Stem cells international 2018, 2018:5416923.

12. Bhatt DK, Chammas R, Daemen T: Resistance Mechanisms Influencing Oncolytic Virotherapy, a Systematic Analysis. 2021, 9(10):1166.

13. Chung SH, Ye XQ, Iwakura Y: Interleukin-17 family members in health and disease. International immunology 2021, 33(12):723–729.

14. Zhao J, Chen X, Herjan T, Li X: The role of interleukin-17 in tumor development and progression. Journal of Experimental Medicine 2019, 217(1).

15. Alizadeh D, Katsanis E, Larmonier N: The multifaceted role of Th17 lymphocytes and their associated cytokines in cancer. Clinical & developmental immunology 2013, 2013:957878.

16. McGeachy MJ, Cua DJ, Gaffen SL: The IL-17 Family of Cytokines in Health and Disease. Immunity 2019, 50(4):892–906.

17. Kuwabara T, Ishikawa F, Kondo M, Kakiuchi T: The Role of IL-17 and Related Cytokines in Inflammatory Autoimmune Diseases. Mediators of inflammation 2017, 2017:3908061.

18. Zhang X, Li B, Lan T, Chiari C, Ye X, Wang K, Chen J: The role of interleukin-17 in inflammation-related cancers. 2025, Volume 15 - 2024.

19. Zhao J, Chen X, Herjan T, Li X: The role of interleukin-17 in tumor development and progression. The Journal of experimental medicine 2020, 217(1).

20. Zhang X, Li B, Lan T, Chiari C, Ye X, Wang K, Chen J: The role of interleukin-17 in inflammation-related cancers. Frontiers in immunology 2024, 15:1479505.

21. Wang S, Wang J, Wu S, Fu Y, Li M: The central role of IL-17 in cancer stemness and immune evasion: A novel axis for overcoming immune checkpoint inhibitor resistance. Critical reviews in oncology/hematology 2026, 217:104999.

22. Burns JS, Manda G: Metabolic Pathways of the Warburg Effect in Health and Disease: Perspectives of Choice, Chain or Chance. International journal of molecular sciences 2017, 18(12).

23. Zhang H, Fan J, Kong D, Sun Y, Zhang Q, Xiang R, Lu S, Yang W, Feng L, Zhang H: Immunometabolism: crosstalk with tumor metabolism and implications for cancer immunotherapy. Molecular cancer 2025, 24(1):249.

24. Kao Y-S, Lauterbach M, Lopez Krol A, Distler U, Godoy GJ, Klein M, Argüello RJ, Boukhallouk F, Vallejo Fuente S, Braband KL et al: Metabolic reprogramming of interleukin-17-producing γδ T cells promotes ACC1-mediated de novo lipogenesis under psoriatic conditions. Nature metabolism 2025, 7(5):966–984.

25. Ma WT, Yao XT, Peng Q, Chen DK: The protective and pathogenic roles of IL-17 in viral infections: friend or foe? Open biology 2019, 9(7):190109.

26. Saddawi-Konefka R, Seelige R, Gross ET, Levy E, Searles SC, Washington A, Jr., Santosa EK, Liu B, O’Sullivan TE, Harismendy O et al: Nrf2 Induces IL-17D to Mediate Tumor and Virus Surveillance. Cell reports 2016, 16(9):2348–2358.

27. Girondel C, Meloche S: Interleukin-17 Receptor D in Physiology, Inflammation and Cancer. 2021, Volume 11 - 2021.

28. Pande S, Yang X, Friesel R: Interleukin-17 receptor D (Sef) is a multi-functional regulator of cell signaling. Cell communication and signaling: CCS 2021, 19(1):6.

29. Ramirez-Carrozzi V, Ota N, Sambandam A, Wong K, Hackney J, Martinez-Martin N, Ouyang W, Pappu R: Cutting Edge: IL-17B Uses IL-17RA and IL-17RB to Induce Type 2 Inflammation from Human Lymphocytes. Journal of immunology (Baltimore, Md: 1950) 2019, 202(7):1935–1941.

30. Bastid J, Dejou C, Docquier A, Bonnefoy N: The Emerging Role of the IL-17B/IL-17RB Pathway in Cancer. 2020, Volume 11 - 2020.

31. El-Tanani M, Rabbani SA, Satyam SM, Rangraze IR, Wali AF, El-Tanani Y, Aljabali AAA: Deciphering the Role of Cancer Stem Cells: Drivers of Tumor Evolution, Therapeutic Resistance, and Precision Medicine Strategies. In: Cancers. vol. 17; 2025: 382.

32. Moitra K: Overcoming Multidrug Resistance in Cancer Stem Cells. BioMed research international 2015, 2015:635745.

33. Zhang Y, Zoltan M, Riquelme E, Xu H, Sahin I, Castro-Pando S, Montiel MF, Chang K, Jiang Z, Ling J et al: Immune Cell Production of Interleukin 17 Induces Stem Cell Features of Pancreatic Intraepithelial Neoplasia Cells. Gastroenterology 2018, 155(1):210–223.e213.

34. Chen XW, Zhou SF: Inflammation, cytokines, the IL-17/IL-6/STAT3/NF-κB axis, and tumorigenesis. Drug design, development and therapy 2015, 9:2941–2946.

35. Eisenreich W, Rudel T, Heesemann J, Goebel W: How Viral and Intracellular Bacterial Pathogens Reprogram the Metabolism of Host Cells to Allow Their Intracellular Replication. 2019, Volume 9 - 2019.

36. Thai M, Graham NA, Braas D, Nehil M, Komisopoulou E, Kurdistani SK, McCormick F, Graeber TG, Christofk HR: Adenovirus E4ORF1-induced MYC activation promotes host cell anabolic glucose metabolism and virus replication. Cell metabolism 2014, 19(4):694–701.

37. Papagiannidis D, Bircham PW, Lüchtenborg C, Pajonk O, Ruffini G, Brügger B, Schuck S: Ice2 promotes ER membrane biogenesis in yeast by inhibiting the conserved lipin phosphatase complex. The EMBO Journal 2021, 40(22):EMBJ2021107958.

38. Sancho P, Barneda D, Heeschen C: Hallmarks of cancer stem cell metabolism. British Journal of Cancer 2016, 114(12):1305–1312.

39. Zaytseva YY, Harris JW, Mitov MI, Kim JT, Butterfield DA, Lee EY, Weiss HL, Gao T, Evers BM: Increased expression of fatty acid synthase provides a survival advantage to colorectal cancer cells via upregulation of cellular respiration. 2015, 6(22).

40. Wold WS: Adenovirus genes that modulate the sensitivity of virus-infected cells to lysis by TNF. Journal of cellular biochemistry 1993, 53(4):329–335.

41. Lukashok SA, Tarassishin L, Li Y, Horwitz MS: An adenovirus inhibitor of tumor necrosis factor alpha-induced apoptosis complexes with dynein and a small GTPase. Journal of virology 2000, 74(10):4705–4709.

42. Choi Y, Bowman JW, Jung JU: Autophagy during viral infection - a double-edged sword. Nature reviews Microbiology 2018, 16(6):341–354.

43. Gao S, Xie X, Fan L, Yu L: Efficacy and safety of IL-17, IL-12/23, and IL-23 inhibitors for psoriatic arthritis: a network meta-analysis of randomized controlled trials. 2025, Volume 16 - 2025.

44. Ungan D, Be C, Baczyk P, Mittermeier S, Lehmann S, Wiesmann C, Huber T, Kolbinger F, Rondeau J-M: IL-17A complexes with therapeutic antibodies exhibit distinct size distributions, potentially contributing to clinically observed immunogenicity. mAbs 2025, 17(1):2575840.

45. Luo Q, Liu Y, Shi K, Shen X, Yang Y, Liang X, Lu L, Qiao W, Chen A, Hong D et al: An autonomous activation of interleukin-17 receptor signaling sustains inflammation and promotes disease progression. Immunity 2023, 56(9):2006–2020.e2006.

